# *Caenorhabditis elegans* Exhibits Positive Gravitaxis

**DOI:** 10.1101/658229

**Authors:** Wei-Long Chen, Hungtang Ko, Han-Sheng Chuang, Haim H. Bau, David Raizen

## Abstract

Whether or not the micro swimmer *Caenorhabditis elegans* senses and respond to gravity is unknown. We find that *C. elegans* aligns its swimming direction with that of the gravity vector (positive gravitaxis). When placed in an aqueous solution that is denser than the animals, they still orient downwards, indicating that non-uniform mass distribution and/or hydrodynamic effects are not responsible for animal’s downward orientation. Paralyzed worms and worms with globally disrupted sensory cilia do not change orientation as they settle in solution, indicating that gravitaxis is an active behavior that requires gravisensation. Other types of sensory driven orientation behaviors cannot explain our observed downward orientation. Like other neural behaviors, the ability to respond to gravity declines with age. Our study establishes gravitaxis in the micro swimmer *C. elegans* and suggests that *C. elegans* can be used as a genetically tractable system to study molecular and neural mechanisms of gravity sensing and orientation.

**Significance Statement:** Understanding how animals respond to gravity is not only of fundamental scientific interest, but has clinical relevance, given the prevalence of postural instability in aged individuals. Determining whether *C. elegans* responds to gravity is important for mechanistic studies of gravity sensing in an experimentally tractable animal, for a better understanding of nematode ecology and evolution, and for studying biological effects of microgravity. Our experiments, which indicate that *C. elegans* senses and responds to gravity, set the stage for mechanistic studies on molecular mechanisms of gravity sensing.

## Introduction

Gravity plays an important role in most life forms on earth, ranging from single cells to plants and animals. Plants’ roots grow in the direction of gravity (positive gravitropism) and shoots grow in the opposite direction (negative gravitropism) to optimize nutrient uptake and exposure to light (1). Aquatic invertebrates use gravity cues to help navigate in the vertical dimension (2). Both terrestrial and aquatic vertebrates know which direction is up. While there are gravity sensory organ differences that relate to the unique ecologies across phylogeny, there are also similarities in the anatomical and physiological principles of such organs. Despite the importance of gravity sensing to life on earth, many molecular components of sensing and responding to gravity remain unknown.

In this study, we examine whether the nematode *Caenorhabditis elegans* (*C. elegans*) senses and responds to the direction of gravity. *C. elegans* offers important experimental advantages, including a small and simple nervous system, accessibility to rapid genetic manipulation and to other powerful experimental tools, and ease of cultivation. There is only one report suggesting that *C. elegans* suspended in solution may orient with the gravitational field (3). Since *C. elegans* is heavier than suspending buffers typically used in laboratories, it settles when suspended in solution (4). Our observations suggest that as wild type animals settle, they also orient their direction of swimming to align with the direction of the gravity vector. Does *C. elegans* sense gravity? Is its response to gravitational forces passive or active? We set to answer these questions in this study.

## Materials and methods

### Worm preparation

On the day prior to the experiment, well-fed fourth larval stage hermaphrodites were placed on an agar plate containing a bacterial lawn of OP50. Day-one adult worms were harvested from the agar plate by floating the worms in M9 buffer and then transferring the worm suspension into a 1.5 mL conical tube. Following centrifugation (4000-5000 rpm) for a few seconds to sediment the worms, the supernatant was decanted. The worms were then washed three times with 1 mL M9 buffer by repeating the centrifugation/decanting steps. We experimented mostly with “recently-fed” worms - the time elapsed from floating the worms off their cultivation plate to the completion of the experiment was < 30 min. A few of the experiments were carried out with “starved” worms - the time elapsed from floating the worms off their cultivation plate to the start of the experiment was ~ 1 hour.

To paralyze wild-type worms, we suspended the worms in one milliliter M9 buffer in a 1.5 mL conical tube and placed them for one hour in a water bath at 40°C. The experiment was then performed at room temperature (21~22°C) within 30 minutes from the removal of the animals from the water bath. Observations of these worms showed that they were fully paralyzed for over 30 minutes after the heat shock

### High density buffer

To achieve a density greater than that of the worms, we mixed a colloidal silica solution (LUDOX HS-40, Sigma, density: 1.3 g/mL at 25°C (5) with M9 buffer at a volume ratio 1:2 to form a solution with density of 1.1 g/mL, which is slightly greater than the worm’s density (~1.07 g/mL (6)). The mixture density was measured directly by weighing one ml of solution. At the density used in our experiment, the suspension behaves like a Newtonian liquid with a viscosity approximately 7 times that of water (7).

### Experimental Apparatus

A cuboid polystyrene cuvette with a square cross-section 12 mm × 12 mm and heights ranging from 45 to 200 mm filled with M9 buffer at room temperature (21~22°C) were used in our settling experiments. 20 μL of a worm suspension at a concentration of about 1.5 worms per microliter was extracted with a plastic tip pipette and transferred into the cuvette by slowly expelling the worms into the cuvette solution either above or just below the liquid surface.

### Imaging

The worms were monitored with two cameras (IMAGINGSOURCE DMK 33GP031 with a 25 mm lens and IMAGINGSOURCE DMK 22BUC03 with a 12 mm lens) acquiring images at 30 frames per second from two orthogonal planes (Fig. 1). One camera focused on the X-Z plane and the other on the Y-Z plane at the cuvette’s center. Each image size was 640 × 480 pixels, which results in an aspect ratio of 4:3. As it settled, a worm stayed about 10 s within the field of view of the two cameras, resulting in about 300 double frames for each worm. Images were processed with a Matlab R2018b graphical user interface (GUI), following the image processing scheme described in (8) and outlined in the Supporting Information (SI-Section S1).

**Figure 1:**
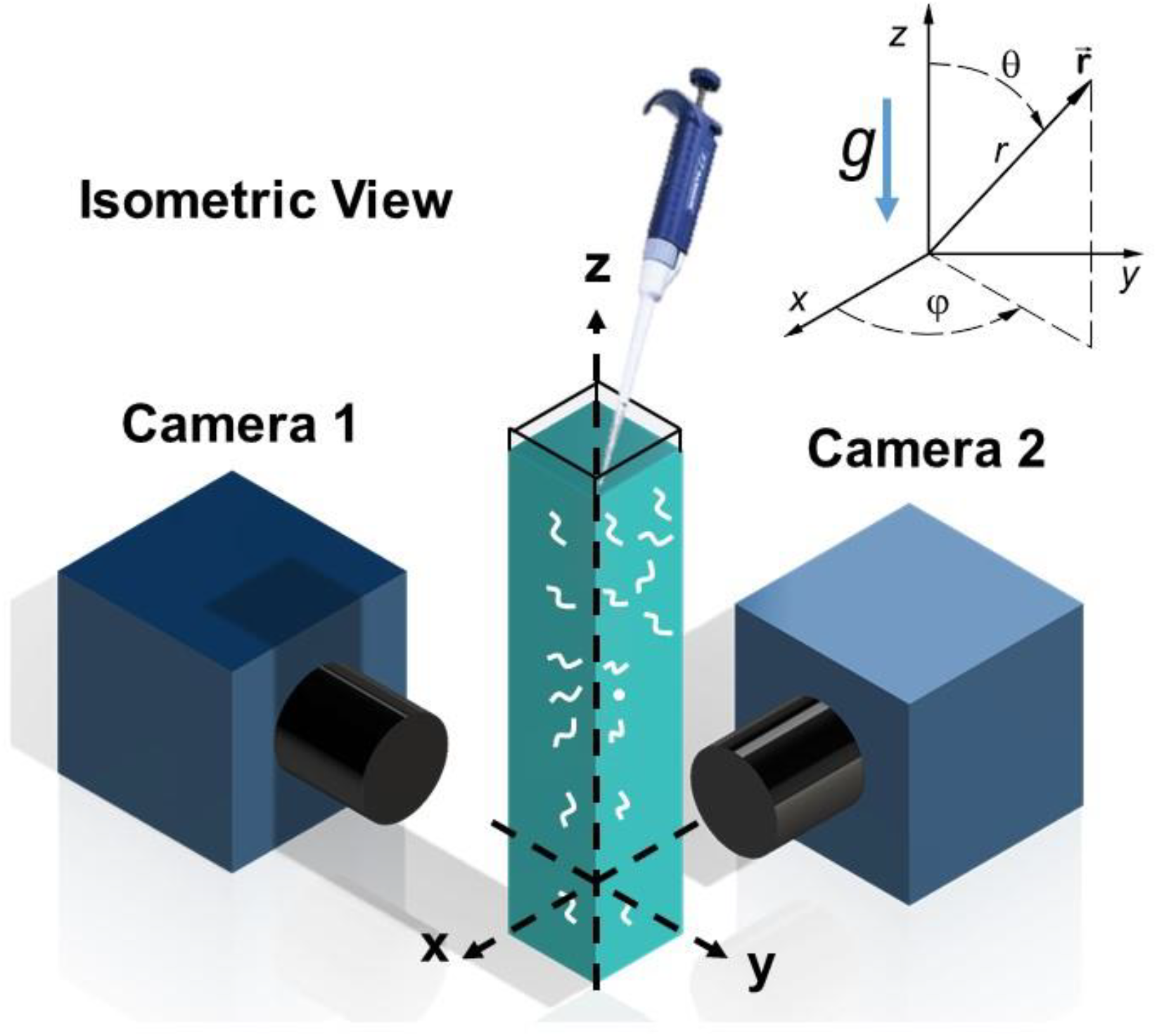
Experimental set-up (Isometric View)

## Results

### Wild Type (WT) *C. elegans* young adults align their swimming direction with the direction of gravity

We inserted first day adult, recently-fed, wild-type (WT) animals just beneath the water surface at the top of our cuvette and monitored the animals’ orientation (*θ*, *ϕ*) as a function of time (Fig. 1). Here, *θ* and *ϕ* are, respectively, the polar (inclination) and azimuthal angles. *θ* = 180° is the direction of gravity. Since the worms (density ~1.07 g/mL) are heavier than water (~1 g/mL), they settled to the cuvette’s bottom.

As time went by, the WT worms varied their swimming direction to align with the direction of the gravity vector. Fig. 2 exemplifies this behavior. The figure depicts time-lapsed frames, 1-second apart, of a WT young adult animal, inserted beneath the water surface and settling in our cuvette. In the first image (A), the animal is a distance ~ 6.5 mm beneath the water surface and faces nearly upwards *θ ~ 5*°. As time increases, the polar angle *θ* gradually increases. In the last frame (J), the animal is ~11.5 mm beneath the water surface and its polar angle *θ* ~ *142*°. The worm has changed its direction of swimming from nearly upwards to nearly downwards. This behavior is exhibited more clearly in panel K, wherein we translated the skeletons of the animal to position their centroids at the same point. Fig. 2K illustrates the animal’s tendency to rotate to align its direction of swimming with the direction of gravity.

**Figure 2:**
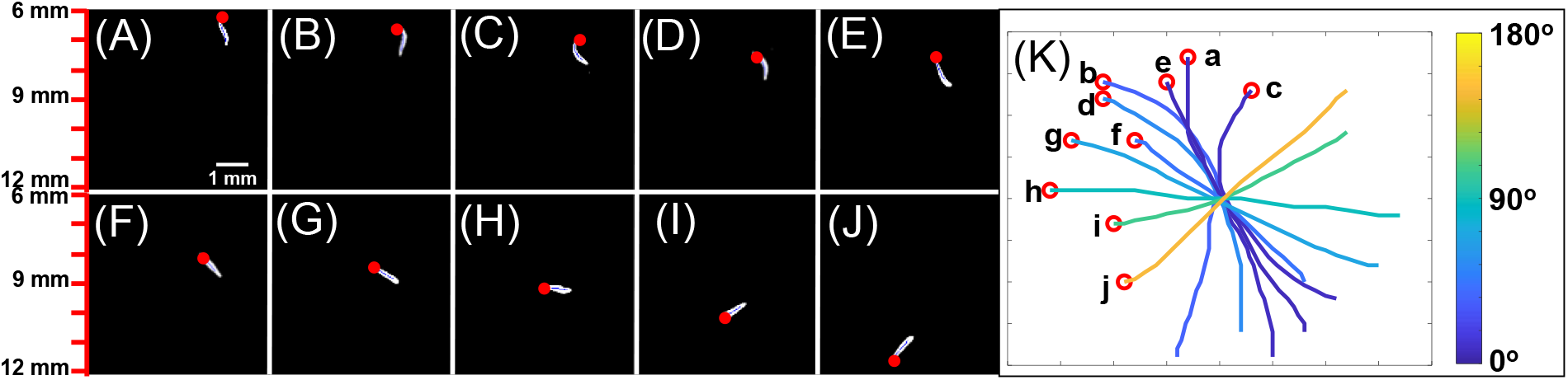
Wild-type animals rotate to align their direction of motion as they descend in solution. (A-J) 10 video frames spaced 1s apart of a descending young adult, wild-type worm. The red dot indicates the position of the worm’s head. The animals are 6-12 mm beneath the water surface. The polar angle varied from 5.1° (frame A) to 141.5 (frame J). (K) The skeletons of the worms from (A-J) were shifted to align their geometric centers to better describe animal’s rotation.

Regardless of initial orientation, given enough time, the animals oriented themselves in the direction of gravity independent of their azimuthal position (SI-Section S2). Fig. 3 depicts the kernel density estimate KDE *f*(*θ*) (an approximation of the probability distribution function, *pdf*) (9) of animals’ orientations at various depths beneath the liquid surface. Close to the liquid’s surface (shortly after release), *f*(*θ*) is nearly symmetric about the horizontal direction (*θ* =*90*°), indicating lack of orientation bias and equal probability towards upward and downward swimming. The KDE resembles a *sin* function that corresponds to a uniform *pdf* in the θ-direction. As time goes by, the KDE function skews in the direction of increasing polar angles, indicating that as the worms descend, they rotate to increase their polar angle and align their direction of swimming with the direction of gravity. Our KDE resembles the von-Mises Fisher directional *pdf* (10) with the mean inclination (polar) angle *θ* = *180*° (SI-Section S4):

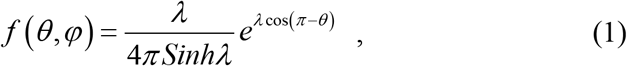

where the *concentration parameter* λ (reciprocal measure of dispersion) is analogous to the inverse of the variance in a normal distribution. *λ* →*0* corresponds to a uniform distribution. Since our data is independent of the azimuthal angle *ϕ*, we integrate in *ϕ* to obtain

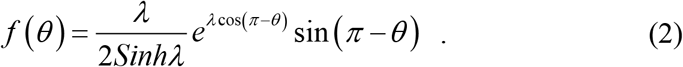

**Figure 3:**
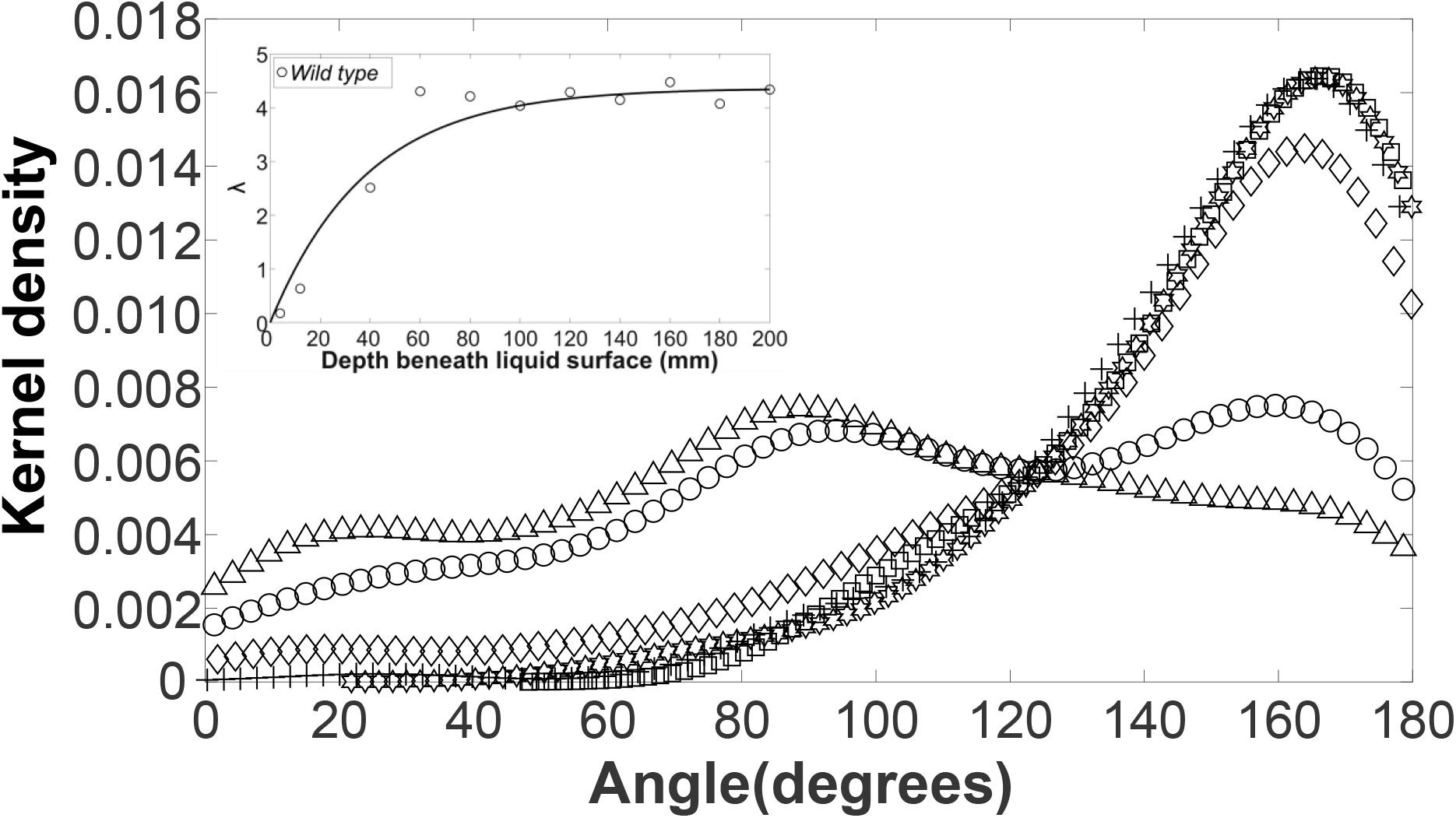
Wild-type worms change their preferred orientation as they settle in solution. Kernel-density of wild-type swimmers’ orientation angle (θ) at positions 4 mm (∆, N=145), 12 mm (○, N=141), 40 mm (◊, N=120), 60 mm(□, N=133), 80 mm (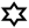, N=126), and 100 mm (+, N=123) beneath the liquid surface. The figure was produced with the Matlab™ function “ksdensity” with “bandwidth” of 15. The inset depicts the concentration parameter λ as a function of the animal’s position (*d* mm) beneath the surface. KDEs for depths >100 mm are provided in SI Section S5.

When λ *≥ 1* and λ *≥ 3*, over 73% and 95% of the animals are oriented, respectively, at a polar angle *θ* > *90*°. We compute the concentration parameter λ for our data by fitting the cumulative distribution function (*cdf*) associated with equation 2 (SI-section S4) to the experimental one. When the animal is at depth *d* = 4 mm beneath the surface *λ* ~ *0.2* (nearly uniform distribution). As the animal’s depth increases (the animal has more time to align with the direction of gravity), the skewness of the KDE and the magnitude of *λ* increase as well. For the well-fed WT animals *λ* increases at the approximate rate of 0.07 per mm of depth until it asymptotes to ~ 4.3 at ~60 mm, and approximately retains this value at depths exceeding 60 mm. KDEs at depths *120 mm* < *d* < 200 *mm* nearly overlap (SI – Section S5). The inset in Fig. 3 depicts the concentration parameter *λ* as a function of the animal’s depth (*d*, mm) beneath the liquid surface. The data is correlated with the expression

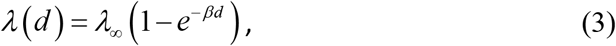

where *λ*_∞_ ~ *4.36* and *β*~ 0.03 mm^−1^. Equation (3) illustrates that after the worm reaches a certain depth, the orientation of the animals attains a stationary state.

The sedimentation velocity of a rigid, cylindrical rod depends on the rod’s orientation with respect to its direction of motion (11); rigid rods settle faster when aligned broadside than when their axis parallels the direction of motion. To test whether this applies to *C. elegans* and to approximately correlate the animal’s depth with its residence time in solution, we examined the worm’s translational and angular velocities. Fig. 4 depicts the velocity (*U*) of young adult WT worms’ centroid in the direction of swimming as a function of (− *cos θ*). The data is scattered along a straight line and correlates well (R^2^=0.89, solid line) with the expression.

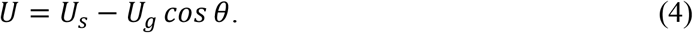

**Figure 4:**
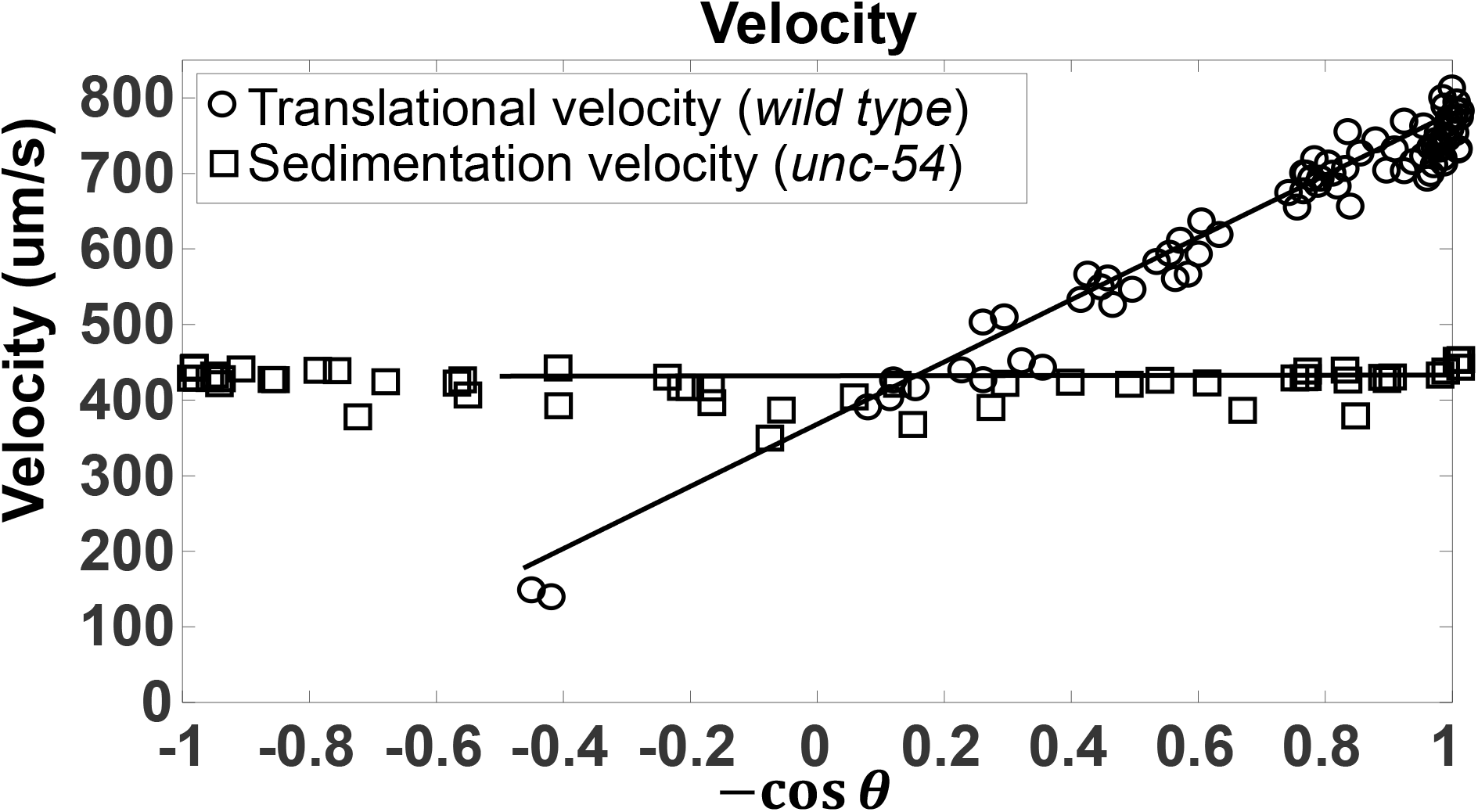
Translational velocity and Sedimentation velocity of worms during gravitaxis. Translational velocity of first day adult WT (N=79) and of motion-impaired adult mutant *unc-54* (N = 52) as functions of -cos θ, where θ =0 corresponds to upward orientation.

We interpret *U*_*s*_ as the animal’s swimming velocity and *U*_*g*_ as the sedimentation velocity in the gravitational field. The term (−*U*_*g*_ *cos θ*) is the projection of the sedimentation velocity on the animal’s swimming direction. In contrast to rigid cylindrical rods, the worm’s settling velocity *U*_*g*_ depends only weakly on orientation (*θ*). This perhaps results from the worm not being perfectly straight and rigid. We estimate 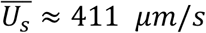 and 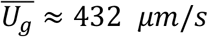. The angular velocity 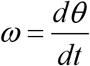 varied widely, but never exceeded *28 degrees/ s*.

To examine reproducibility of our data, we repeated our experiments in two continents (USA and Taiwan) and a few weeks apart and obtained similar results (e.g., **SI - Fig. S3**). We also demonstrated that the animals’ azimuthal angle *ϕ* was uniformly distributed (**SI - Fig. S4**), suggesting that convective currents in our apparatus, if any, are unlikely to have biased our data.

In summary, WT animals align their swimming direction with the gravity vector. Can this alignment be attributed to non-uniform mass distribution along the animal’s length?

### Paralyzed WT animals do not align with the direction of the gravity vector

A plausible cause for animals to align with the direction of gravity is a non-uniform mass distribution along the animal’s body. Animals store fat primarily in the intestine (12), which is not present in the anterior 1/5^th^ of the worm’s body; therefore, the worm may be head-heavy. If significant, a non-uniform mass distribution would cause animals to rotate in a gravitational field. To isolate potential effects of non-uniform mass distribution, we experimented with paralyzed animals.

We achieved muscle paralysis by either exposing WT animals to high temperature (heat-shock) or by testing animals that carry a mutation in the major muscle myosin gene *unc-54* (13). Both heat-shocked WT animals and *unc-54* mutants maintained their initial inclination (polar) angle and did not align with the direction of gravity as they descended. Their settling velocity (*U*_*g*_ ~ 432 *μm/s*) is in excellent agreement with the estimated contribution of gravitational settling to the velocity of WT animals (equation 3 and Fig. 4).

Both paralyzed WT and *unc-54* did not show any orientation preference during sedimentation. The KDEs of the heat-shocked WT (**SI-Fig. S10**, 4 mm < d < 100 mm, and Fig. 5, 120 mm < d < 200 mm) and *unc-54* (**Fig. S11**, d = 40 mm, which is sufficiently far beneath the water surface to allow animals to begin to adjust their polar angle) resemble a uniform distribution in the polar angle and their descent angle *θ* is nearly symmetric with respect to θ~90°. Likewise, λ of paralyzed WT animals ranged from 0.01 to 0.33 consistent with a nearly uniform distribution (inset in Fig. 5). Contrast Figs. 5 and **S10** with Figs. 3 and **S9**. The difference is striking. Active WT animals rotate to align with the gravity vector while paralyzed animals do not vary their polar angle θ during their descent. The kernel distribution estimates of Figs. 5 and **S10** are statistically distinct from the kernel distribution estimate of active animals Fig. 3 and **S9** (p < 0.0001, Mann Whitney test (9)).

**Figure 5:**
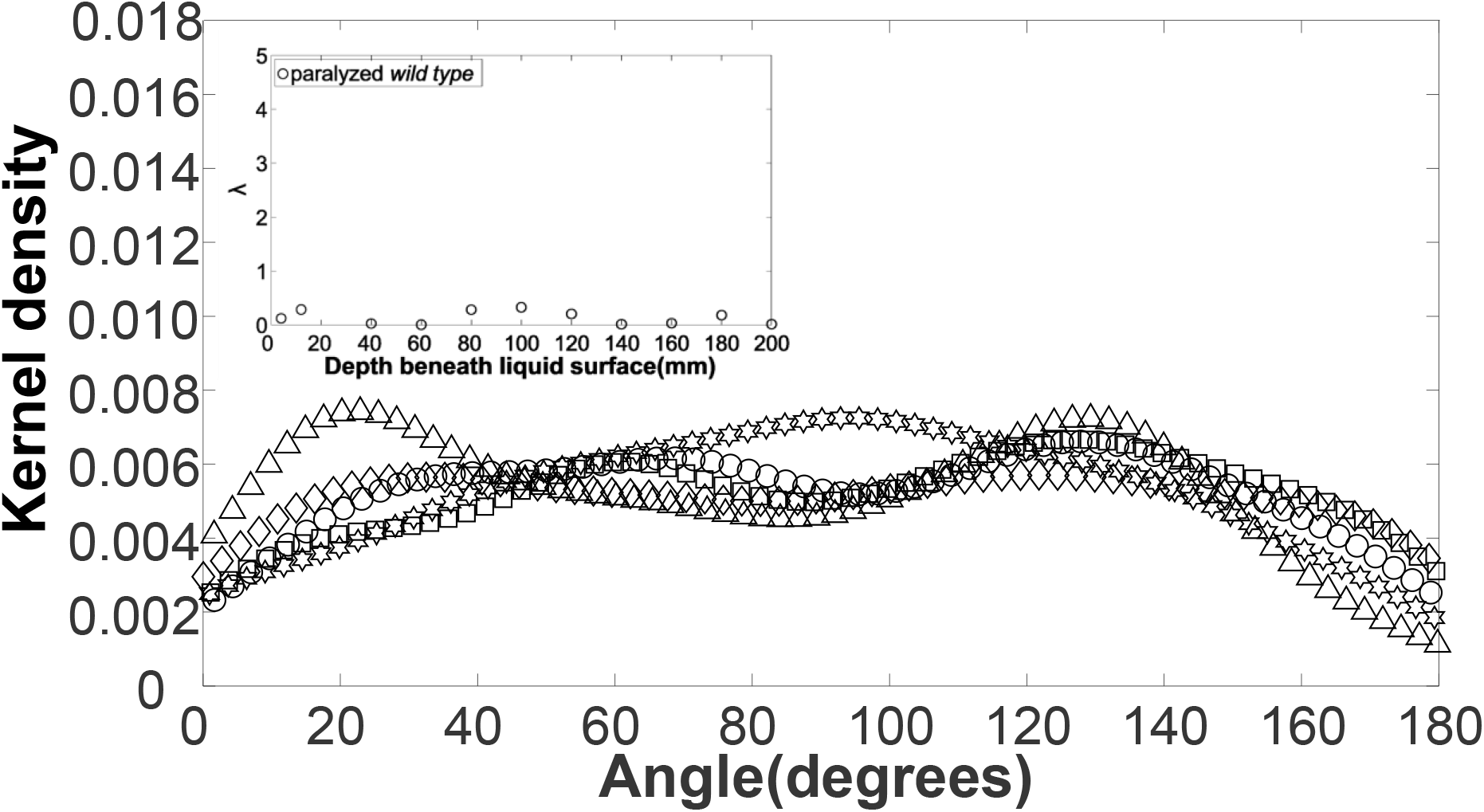
Paralyzed WT worms retain random distribution of their orientation as they settle in solution. Kernel (probability) density estimate (KDE) of heat-shocked paralyzed WT animals at positions 120 mm (∆, N=125), 140 mm (○, N=128), 160 mm (◊, N=133), 180 mm (□, N=127), and 200 mm (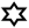, N=126) beneath the liquid surface. The bandwidth of the KDE smoothing window is 15. See SI for KDEs of paralyzed WT at smaller depths (**Fig. S10**). Inset: the concentration parameter λ as a function of depth. λ remains close to zero consistent with uniform (random) distribution.

In summary, the marked difference between active and paralyzed animals indicates that the propensity to align with the gravity vector is mediated by active mechanisms. These experiments lend further support to our earlier conclusion that factors such as convective currents in our cuvette, if any, are not responsible for animal’s orientation since they would have similarly impacted active and inactive animals. Does the level of animal’s activity affect how fast it orients itself with the gravitational field?

### Starved animals and animals defective in muscle function (*unc-29*) align with the direction of the gravity vector at a slower rate than well-fed WT animals

To determine whether the propensity for positive gravitaxis behavior is affected by the dietary history or by mild impairment (non-paralysis) in body movements, we experimented with starved (> 1 hour from last feeding) WT animals (**Figs. S13** and **S14)** and with *unc-29* mutants, which are mildly defective in muscle function due to a mutation in an acetylcholine receptor subunit (14) (**SI Figs. S15** and **S16**). In both cases, as the animals’ depth beneath the liquid surface (and residence time) increased, so did the skewness of their KDEs and the magnitude of their concentration factor λ (**SI Fig. S17**), indicating that these animals still align with the direction of the gravity vector; albeit at a slower rate than the well-fed, WT animals. The concentration factor λ of the starved WT animals increased at the approximate rate of 0.03 mm^−1^ with depth, about half that of the well-fed animals, until it attained the nearly stationary value of 3.2 at 100 mm and greater depths. The concentration factor λ of *unc-29* mutants increased at the approximate rate of 0.007 mm^−1^ and attained *λ* ~ *1.5* at the depth of 200 mm, which is the largest depth available in our experimental apparatus. Although the concentration factor λ of *unc-29* mutants did not show fully saturation, it did increase with time, in clear contrast to fully paralyzed mutants.

In summary, the rate of animal’s alignment with the direction of the gravitational field declines as the animal’s swimming vigor decreases. Importantly, even animals with reduced swimming vigor, when given enough time, orient to align with the gravity vector. Our observations suggest that positive gravitaxis behavior does not solely rely on vigorous muscle movements. Can the propensity to align with the direction of gravity be attributed to hydrodynamic effects?

### Gravitaxis does not result from hydrodynamic effects

We reasoned that if the downward swimming orientation were the result of interactions between the flow field induced by the swimmer and the flow field associated with downward sedimentation than, based on symmetry arguments, animals sedimenting *upward* should align with the direction that is opposite to the direction to the gravity vector. To test this hypothesis, we suspended well-fed WT animals beneath the surface of a LUDOX suspension that has density slightly greater than that of the animals. Our experiments were complicated by the animals floating to the surface and the LUDOX suspension having viscosity greater than water, decreasing the rotational velocity of the animals and allowing them less time to orient in the gravitational field. In contrast to the predictions based on hydrodynamic symmetry considerations, the animals rotated to orient *downwards* and swim in the direction of the gravity vector. Fig. 6 (A-J) shows 10 video frames, spaced 1s apart, of a young adult, well-fed, WT worm. The red dot indicates the position of the worm’s head. In the 10-seconds period of observation, the polar angle increased from an initial value of *49.1*° in frame A to a value of *138.9*° in frame J. Frame K depicts the skeletons of the worms from frames (A-J) shifted to align their geometric centers.

**Figure 6:**
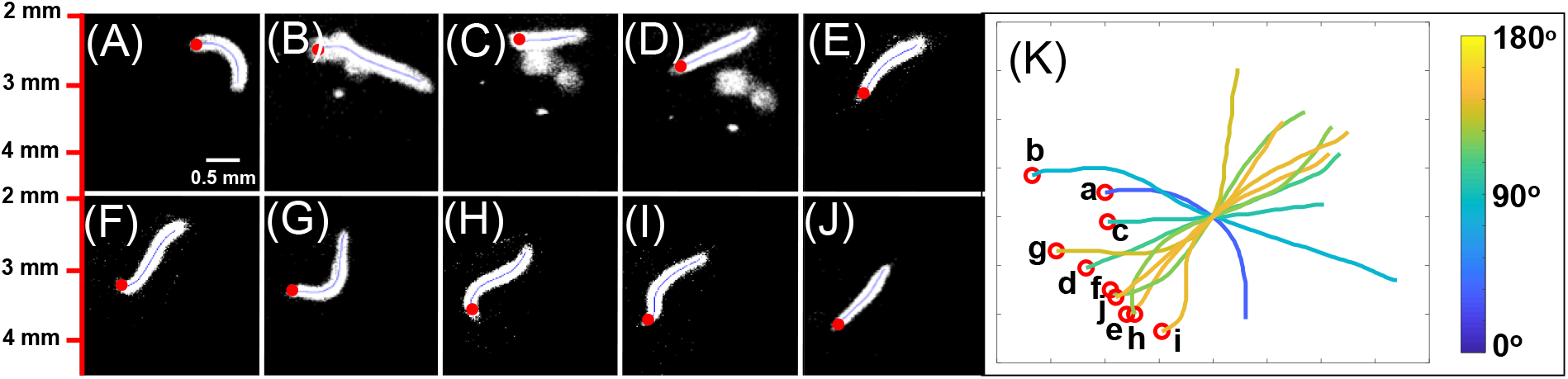
Wild-type animals rotate to align their direction of motion downward when suspended in a solution denser than the animals. (A-J) 10 video frames spaced 1s apart of a young adult, wild-type worm. The red dot indicates the position of the worm’s head. The polar angle varied from 49.1° in frame A to 138.9° in frame J. (**K**) The skeletons of the worms in (A-J) were shifted to align their geometric centers.

Fig. 7 depicts the kernel (probability) density estimate of the polar angle *θ* shortly after the animals’ introduction into the suspension, 5 s later, and 10 s later. At short times, the KDE resembles a *sin* function, characteristic of a uniform *pdf* in *θ*. As time passes, the peak of the kernel density estimate shifts to larger values of the angle *θ* and the magnitude of the concentration factor *λ* increases from nearly zero to 2.7. Therefore, animals suspended in a liquid that is either lighter (Fig. 3) or heavier (Fig. 7) than themselves rotate to swim in the direction of the gravity vector. In summary, our observation of downward swimming even when sedimenting upwards indicates that hydrodynamics are not responsible for animals’ orientation. Gravitaxis is not caused by hydrodynamic effects.

**Figure 7:**
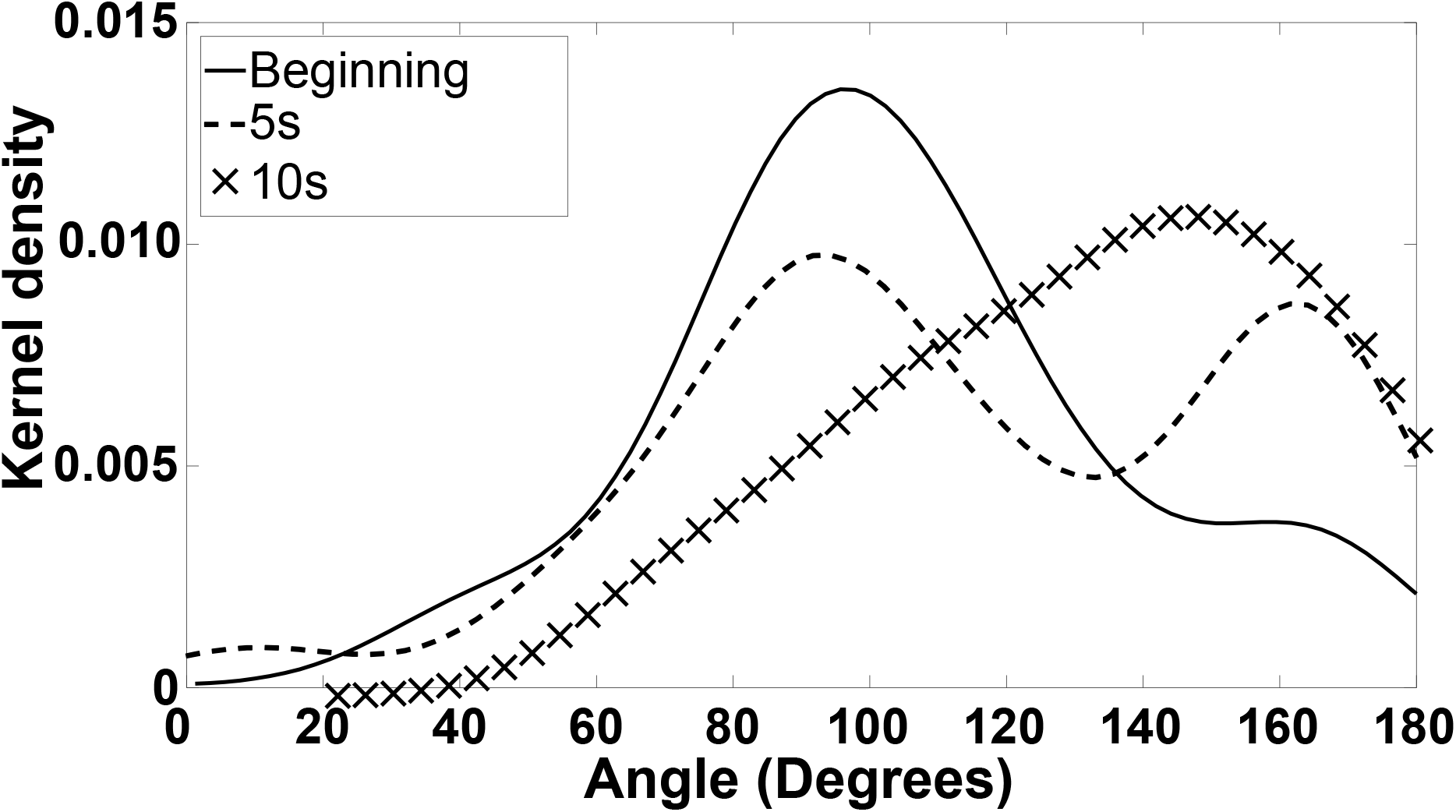
Gravitaxis does not affected by direction of buoyancy. The kernel (probability) density estimate of orientation angle of animals suspended in LUDOX HS-40 suspension (density 1.1 g/mL and viscosity about 7 times that of water) shortly (< 2s) after the animals introduction into the suspension, 5s later, and 10s later. N_0_ = 31, N_5s_ = 30, and N_10s_ = 36. In depicting the KDE curves, we used Matlab™ default values.

There is another important conclusion that we can draw from this experiment. In both vertebrates and some aquatic invertebrates, gravity is sensed via the interaction between a proof mass (a mass denser than its surrounding) and sensory cilia of specialized cells internal to the animal. In contrast, certain insects, such as strepsiptera species, use their head as a proof mass to sense gravity. As the animal’s head falls, the head brushes against hairs between the head and thorax (2). By monitoring the direction of hair deformation, the animal senses the direction of gravity. Our experiment indicates that the gravity sensing mechanisms in *C. elegans* are interior to the animal’s body, as is the case in vertebrates and some aquatic invertebrates and are not affected by the direction of buoyancy.

### Magnetotaxis and other taxis are not responsible for the worm orientation in gravitational field

Vidal-Gadea et al (15) report that *C, elegans* orients to the earth’s magnetic field during vertical burrowing migrations. Well-fed adult worms of the N2 Bristol strain, which was isolated in the Northern Hemisphere, migrated up, while starved N2 worms migrated down. In contrast, well-fed adult worms of the AB1 Adelaide strain, which was isolated in the Southern Hemisphere, migrated down while starved AB1 worms migrated up in response to the same magnetic field. We have not observed similar tendencies in our experiments with *C. elegans* in solution. In our experiments, both well-fed (Figs. 2 and 3) and starved (SI Figs. S13, S14, and S17) WT animals oriented with the direction of the gravity field and swam downwards. Well-fed AB1 worms (SI **Figs. S18 – S20**), like well-fed N2 worms, oriented downwards in the vertical liquid column. Hence, the taxis mechanisms identified in reference (15) are unlikely to explain our observations.

Since our experiments took place in a vessel subjected to uniform room light and temperature, there is no gradient of light intensity or temperature and therefore no phototaxis or thermotaxic stimulation. While *C. elegans* prefers low oxygen tensions (16, 17), our observations are unlikely to be explained by aerotaxis behavior because the concentrations of gases are nearly uniform in our aqueous column, aside from O_2_ consumption and C0_2_ production by the worms, which is likely to be negligible on the time scale of our experiment. Moreover, whereas in a low water density solution, the animals sedimented downward and aggregated at the bottom of the water column, in high-density solution, the animals sedimented upwards and congregated at the top of the water column. Any gaseous gradient caused by consumption of oxygen or generation of carbon dioxide by aggregated animals would be reversed in the high-density solution. Our observation that animal orientated downward regardless of where the animals aggregated (at the top or the bottom) further indicates that gaseous gradients do not explain our observations. In conclusion, given our exclusion of other known taxis behaviors, our observations indicate that the animals respond to gravity when swimming in an aqueous solution.

### Gravity sensing and ability to orient in gravitational field declines with age

Many of *C. elegans* sensing capabilities deteriorate with age (18). Some of this deterioration is identifiable with specific neuronal deficits. For example, the sensory dendrites of the FLP and PVD neurons, which are required for response to certain mechanical stimuli, degenerate with age (19). To examine aging worms’ ability to sense and orient in gravitational field, we measured the angle of descent as a function of animal’s age (Fig. 8) when located 40 mm beneath the liquid surface. The aged animal (day 6) has a broader kernel density estimate than the young adult (day 1), indicating that a greater fraction of the animals failed to align with the direction of gravity. The concentration parameter λ (inset in Fig. 8) decreases as the animal’s age increases. Day 1 and day 2 adults animals behaved similarly with λ = 2.9 (N = 87) and 3.1 (N = 62), respectively. After day 2 of adulthood, there was a gradual decline in λ. At day 6 of adulthood, which corresponds to a mid-life aged animal, λ ~ 1.4 (N = 50), which is significantly smaller than that of day 1 adults (p = 0.0012, Mann Whitney test). In the age range between 1 and 5 days, there is a gradual decline in the animals’ ability to react to gravitational field.

**Figure 8:**
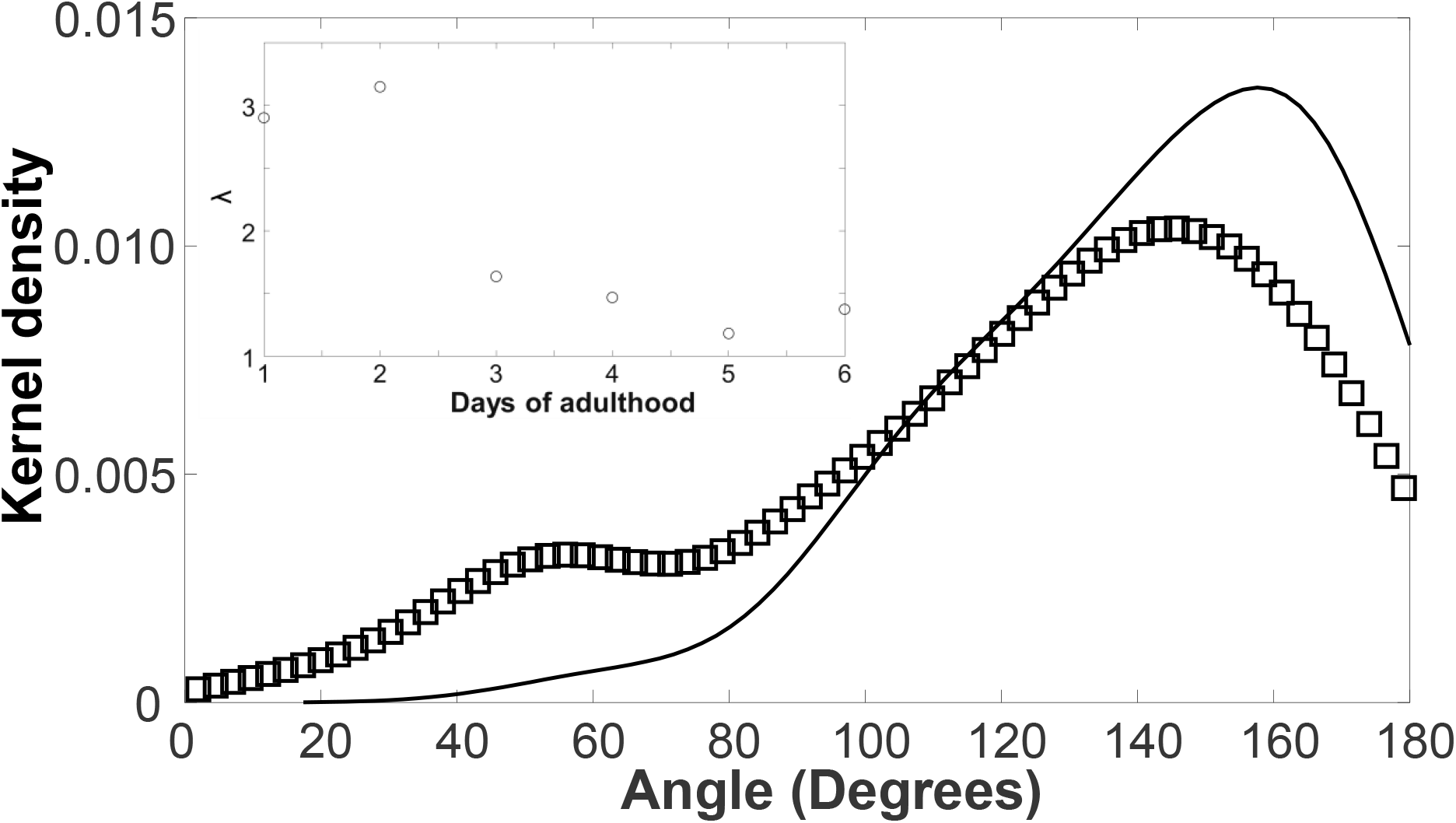
Aging adults have impaired ability to orient with the gravity vector. Kernel-(probability) density estimate of the orientation angle *θ* of day 1 (AD 1, solid line) and day 6 (AD6, □) adults at 40 mm beneath liquid surface. The inset depicts the concentration parameter λ as a function of age. For day 6 animals, p < 0.01 (**, Mann Whitney Test). N_AD1_ = 87, N_AD2_ = 62, N_AD3_ = 60, N_AD4_ = 55, N_AD5_ = 40, N_AD6_ = 50, In depicting the KDE curves, we used Matlab™ default values.

### Gravitaxis requires sensory neurons function

We reasoned that if the worm senses gravity and deliberately orients in the downward direction, we should be able to impair this behavior by selectively disrupting sensation with minimal impairment of movement. Many sensory functions of *C. elegans* such as olfaction, gustation, thermosensation, nose-touch, magnetoreception, and electrosensation are mediated by neurons that extend cilia to the nose of the animal. We hypothesize that gravity sensation too is mediated by ciliated sensory neurons. To test this hypothesis, we analyzed the angle of descent of animals mutant for the gene *osm-6* or for the gene *che-2*, which encode, respectively, intra flagellar protein 52 (IFT52) and IFT80 and in which sensory cilia are globally disrupted (20).

The polar angles of the mutants *che-2* and *osm-6* did not vary as they descended. The kernel-density estimate plots (Fig. 9, 40 mm beneath liquid surface) suggest that both *che-2* and *osm-6* retain nearly uniform kernel density estimates with λ = 0.1 (N = 51) and 0.6 (N = 70), respectively. Their kernel density estimates were statistically distinct from the WT control (p < 0.0001, Mann Whitney Test). Our data suggests that the cilia-mutant animals show little preference in their angle of descent. We conclude that ciliated sensory neurons are necessary for gravitaxis.

**Figure 9:**
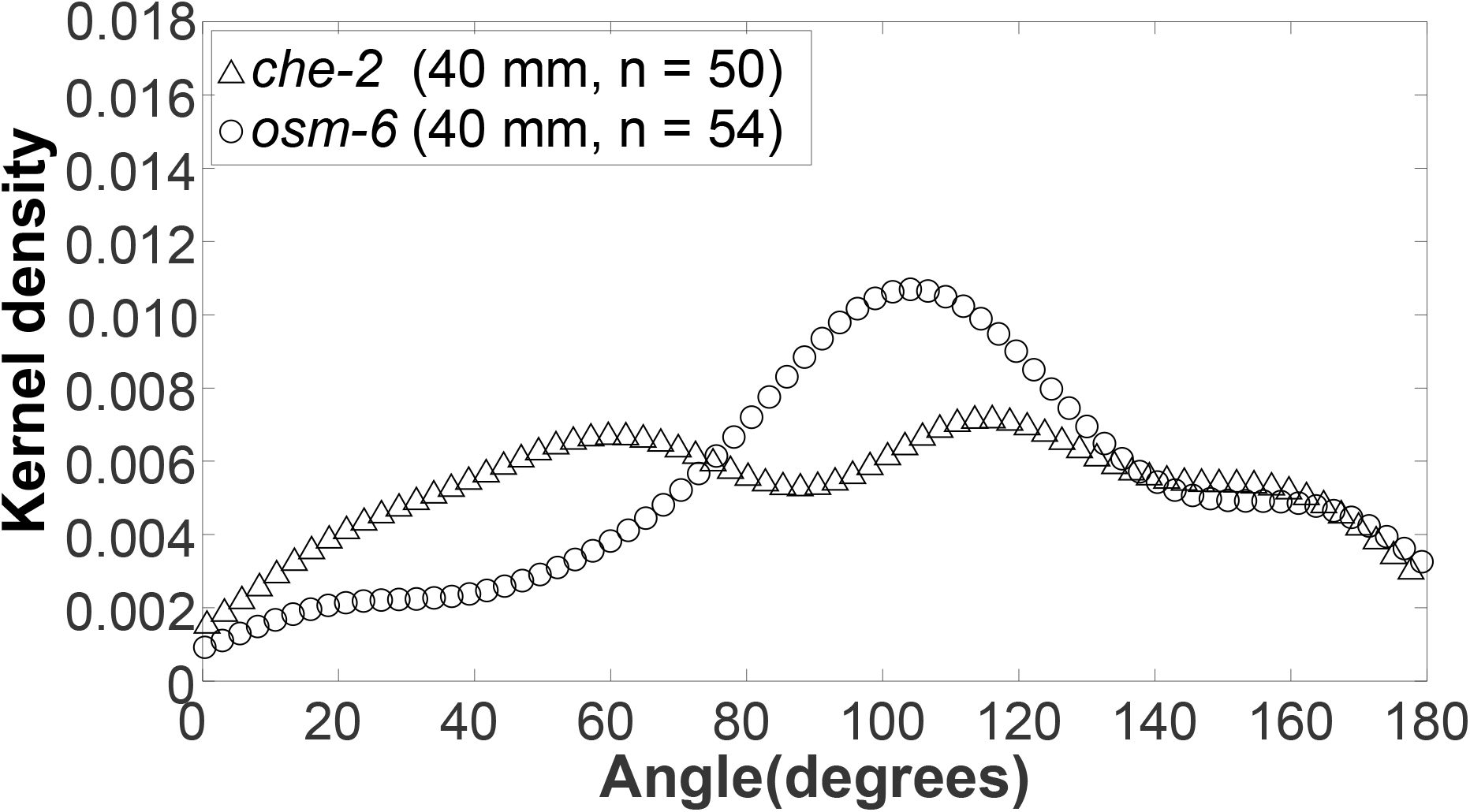
Sensory mutants *che-2* and *osm-6* show defects in downward orientation. Kernel-density estimate plot of angle of descent of sensory mutants and of wild-type controls at 40 mm beneath liquid surface. The distributions of angles of descent of *che-2* and *osm-6* mutants are all broader than that of wild-type animals and approximate random distribution. Compared to WT distribution, p < 0.0001 (Mann Whitney Test). N_WT_=87, N_che-2_=51, and N_osm-6_=70. In depicting the KDE curves, we used Matlab™ default values.

## Discussion

Hydrodynamists characterize motion of objects in fluids based on the relative importance of inertial and viscous forces, quantified by the Reynolds number *Re*=*U D*/*ν*. Where U is the object velocity, *D* is the object’s characteristic length, and *ν* is the kinematic viscosity of the suspending fluid. The combined swimming and settling velocity *U* of *C. elegans* is less than *1 mm/s*. The characteristic length scale, *e.g.*, the diameter of *C. elegans*, *D~80* μm and the kinematic viscosity of water *ν*~*10*^*−6*^ m^2^/s. In all our experiments, *Re<0.1*, viscous effects dominate, and the equations of motion are linear (Stokes equation). When inertia effects are negligible (*Re→0*), cylindrical objects with fore–aft symmetry released with an inclination angle *θ* sediment, in an unbounded medium, with the angle of release (11). Objects with non-uniform mass distribution turn to bring their center of mass beneath their centroid. If *C. elegans* were head heavy, it would eventually descend at *θ =180°* when *Re→0*. In the presence of weak inertia (*Re*>*0*), a cylindrical object turns to attain horizontal (broadside, *θ =90°*) posture and then settles horizontally with its center of mass descending in the direction of gravity (21).

In our experiments, paralyzed WT animals settling in water retained their initial orientation (polar) angle and did not show any tendency to align with the direction of the gravity vector, consistent with low Reynolds number hydrodynamic theory for cylindrical objects with fore– aft symmetry. This suggests that non-uniform mass distribution, if any, along the animal’s length is insignificant. Additionally, our data indicates lack of convective currents in the experimental apparatus that might have affected animals’ orientation. Since motion-impaired animals do not align with the direction of the gravity vector, such alignment requires active mechanisms.

Prior studies indicate that *C. elegans* traits such as gait-synchronization (22), tendency to swim against the flow (rheotaxis) (23, 24), and tendency to swim along surfaces (bordertaxis) (25) are involuntary and can be explained by simple mechanics. Could an interaction between the flow-field induced by the animal’s swimming gait and the flow-field induced by the animal’s sedimentation cause the animal to rotate and align with the gravity field? If such an alignment mechanism existed, one would conclude, based on symmetry arguments, that animals suspended in a liquid denser than them would align in the opposite direction to that of the gravity vector. Our experiments with WT animals in LUDOX solution that is denser than the animals (Figs. 6 and 7) indicate that this is not the case. Hence, we exclude hydrodynamics as a possible explanation for gravitaxis.

Prior workers (15) reported that well-fed N2 strain (isolated in the Northern Hemisphere) and the well-fed AB1 strain (isolated in Southern Hemisphere) crawl in opposite directions in response to the same magnetic field. Moreover, starved N2 worms crawl in the opposite direction to that of well-fed N2 worms in a magnetic field. In contrast, we observed that both N2 and AB1 worms suspended in solution, regardless of being well-fed or starved, oriented downwards in our vertical liquid column, demonstrating that mechanisms identified in reference (15) do <u>not</u> explain our observations.

Our observations that *che-2* and *osm-6* mutants, which have the necessary motility to align with the direction of gravity, fail to do so, further support our conclusion that gravitaxis in *C. elegans* is deliberate, resulting from the animal’s ability to sense the direction of gravity and act on this information.

Gravity-sensing organs in invertebrates may be either external or internal to the animal’s body. The organ responsible for gravity sensing in *C. elegans* is still elusive. While a proof mass similar to that seen in vertebrate inner ears has not been reported in ultrastructural studies of *C. elegans*, it is possible that such a mass may have escaped detection due to its destruction during tissue fixation and preparation processes.

WT *C. elegans* exhibits positive gravitaxis in both suspending medium that is lighter and a suspending medium that is denser than the animal, demonstrating that gravity perception is not affected by the density of the medium external to the animal. Therefore, the organ responsible for graviatxis in *C. elegans* must be internal.

The evolutionary causes of positive gravitaxis behavior in *C. elegans* are a subject of speculation. One possibility is that when dwelling in wet soil, downward migration would keep the worms moist and away from the drying surface as well as distance them from air-borne predators. Downward migration in bodies of water, may provide protection from predators as well as direct the worms towards sources of food such as underwater flora and associated bacteria.

Regardless of the reason for gravitaxis, we have here shown that the microscopic nematode *C. elegans* orients its swimming direction to align with the direction of the gravity vector, and that this behavior is not the result of an unequal distribution of mass, hydrodynamic interactions, experimental artifacts, and other types of sensory-driven movements, or hydrodynamic interactions. Taken together, our results indicate that *C. elegans* can sense and respond to the force of gravity. Our results suggest the possibility of leveraging the powerful genetic and physiological toolkit of *C. elegans* to elucidate the molecular and circuit mechanisms for gravity sensing – mechanisms that are still elusive.

## Acknowledgements

We thank Drs. Christopher Fang-Yen and Abraham Wyner for useful discussions. W.L.C was funded by the Ministry of Education of Taiwan, Global Networking Talent 3.0 Plan (GNT3.0), and the Medical Device Innovation Center, National Cheng Kung University.

## Authors Contributions

DR and HHB planned experiments, HK carried out preliminary experiments, and WLC performed the majority of experiments in the USA and in Taiwan, the latter with supervision by HSC. DR, HHB, and WLC wrote the paper. All authors read and approved the paper,

