## Supplementary material for "*Caenorhabditis elegans* Exhibits Positive Gravitaxis": SI

### Supplemental Information

Wei-Long Chen[1-3], Hungtang Ko[1], Han-Sheng Chuang[3] Haim H. Bau[1], and David Raizen[2]

1. Dept. Mechanical Engineering and Applied Mechanics, University of Pennsylvania, Philadelphia, PA
2. Dept. of Neurology, Perelman School of Medicine, University of Pennsylvania, Philadelphia, PA
3. Current address: Department of Biomedical Engineering, National Cheng Kung University (NCKU), Taiwan

**S1. Supplementary Materials and methods:**

**Strains:** *C. elegans* strains used in this study were: N2 (*wild-type,* WT), AB1 (wild-isolate from Australia), CB190 (*unc-54(e190)*), CB1072 (*unc-29(e1072)),* CB1033 (*che-2(e1033)*), and PR811 (*osm-6(p811)*). The strains were obtained from the Caenorhabditis Genetics Center (CGC). Worms were cultivated in a 20ºC incubator on the surface of NGM agar plate (5.5 cm diameter; 11 mL total volume) seeded with *Escherichia coli OP50* as the food source. All experiments were performed with hermaphrodites.

**Image processing:** Images were processed with a Matlab R2018b Graphical User Interface (GUI), following the image processing scheme described in (16) and outlined below.

**Data analysis:** We denote the coordinates (**Fig. 1**) in the X-Z plane as $(x_{nXZ}, z_{nXY})$ and in the Y-Z plane as $(y_{nZY}, z_{nZY})$, where *n* is the frame number. The cameras' pixel size was correlated with the physical length by placing a ruler in front of the camera at the same distance from the camera lens as the cuvette's mid-plane. The pixel size P in the XZ plane was $P_{XZ} = 18.8\ \mu m/pixel$, and in the Y plane was $P_{YZ} = 22.0\ \mu m/pixel$. Accordingly, the distances

Rx, Ry, and Rz between the animal's head (H) and tail (T) along Cartesian coordinates were: $R_{n,x}^{TH} = P_{XZ}(x_{n,XZ}^{T} - x_{n,XZ}^{H})$ ; $R_{n,z}^{TH} = P_{XZ}(y_{n,XZ}^{T} - y_{n,XZ}^{H}) = P_{YZ}(y_{n,YZ}^{T} - y_{n,YZ}^{H})$ ; and $R_{n,z}^{TH} = P_{YZ}(z_{n,YZ}^{T} - z_{n,YZ}^{H})$. The distance between the animal's head and tail

$$||\overline{TH_n}|| = \sqrt{{R_{n,x}^{TH}}^2 + {R_{n,y}^{TH}}^2 + {R_{n,z}^{TH}}^2} \quad \text{(S1)}$$

The inclination (polar) angle with respect to the vertical axis

$$\theta_n = \cos^{-1} \frac{R_{n,z}^{TH}}{||\overline{TH_n}||}. \quad \text{(S2)}$$

The angle $\theta = 0°$ corresponds to the upward direction. The azimuthal angle

$$\varphi_n = \cos^{-1} \frac{R_{n,x}^{TH}}{\sqrt{{R_{n,x}^{TH}}^2 + {R_{n,y}^{TH}}^2}}. \quad \text{(S3)}$$

The algorithm was verified by reproducing the dimensions and inclination angle of a 3D-printed calibration jig (Supplement S2).

To monitor the worms' orientation as a function of residence time in solution, images of worms were recorded and processed at various positions beneath the liquid surface (**Fig. 1**). The fields of view of the cameras covered 640 × 480 pixels, which corresponds to a vertical distance of approximately 9 mm.

The position of the worm's centroid (geometric center)

$x_{n,XZ}^{C} = \frac{(x_{n,XZ}^{H} + x_{n,XZ}^{T})}{2}$; $y_{n,XZ}^{C} = \frac{(y_{n,XZ}^{H} + y_{n,XZ}^{T})}{2} = \frac{(y_{n,YZ}^{H} + y_{n,YZ}^{T})}{2}$; and $z_{n,YZ}^{C} = \frac{(z_{n,YZ}^{H} + z_{n,YZ}^{T})}{2}$.

The components of the worm's centroid displacement between subsequent frames is given by $\Delta R_{n,x}^{C} = P_{XZ}(x_{n+1,XZ}^{C} - x_{n,XZ}^{C})$; $\Delta R_{n,z}^{C} = P_{XZ}\left(y_{n+1,XZ}^{C} - y_{n,XZ}^{C}\right) = P_{YZ}\left(y_{n+1,YZ}^{C} - y_{n,YZ}^{C}\right)$; and $\Delta R_{n,y}^{C} = P_{ZY}(z_{n+1,YZ}^{C} - z_{n,YZ}^{C})$. The displacement of the worm's centroid between frames

$$\Delta D_n = \sqrt{{\Delta R_{n,x}^{C}}^2 + {\Delta R_{n,y}^{C}}^2 + {\Delta R_{n,z}^{C}}^2}. \quad \text{(S4)}$$

The velocity of the worm is

$$U_n = \Delta D_n \times 30 \ \ (\mu m/s), \tag{S5}$$

where the factor 30 reflects the video frame rate. The polar angular velocity

$$\omega_n = \langle \theta_{n+1} - \theta_n \rangle \times 30 \ \ (rad/s). \tag{S6}$$

**Image Processing Algorithm**

Images were processed with Matlab R2018b GUI, following the image processing scheme describe in (8). Briefly,

1. The grayscale threshold for detecting worms was manually adjusted using the ImageJ program to transform the captured grayscale images into binary black/white images. The gray scale threshold for detecting a worm varied between experiments and ranged from 32 to 85.
2. Background Subtraction. In the first frame (n=0) of each video, we manually defined an imaging region. The region outside this imaging region was assigned a binary scale zero (black). The mode of the binary scale of the imaging region, excluding the worm, was assigned to the region occupied by the worm. The resulting image served as the background frame and was subtracted from all frames in the video.
3. A tight bounding box containing the worm was manually defined in the first frame. The positions of the tips of worm's head and tail were manually marked. We then defined in frame (n+1) an extended bounding box with 10 pixels added to the width and length of the bounding box of frame n to form a search-region for the worm following its displacement in the time span between frame n and frame ($n+1$).
4. To reduce noise, each frame was filtered by replacing each pixel's value with the average

of its 9 neighbors found in a 3 pixel x 3 pixel surrounding square.

5. Next, we used Matlab's edge detection function CANNY to locate the worm's boundary. The contour was smoothed with Matlab's functions "strel", "imdilate", "imerode", and "imfill".

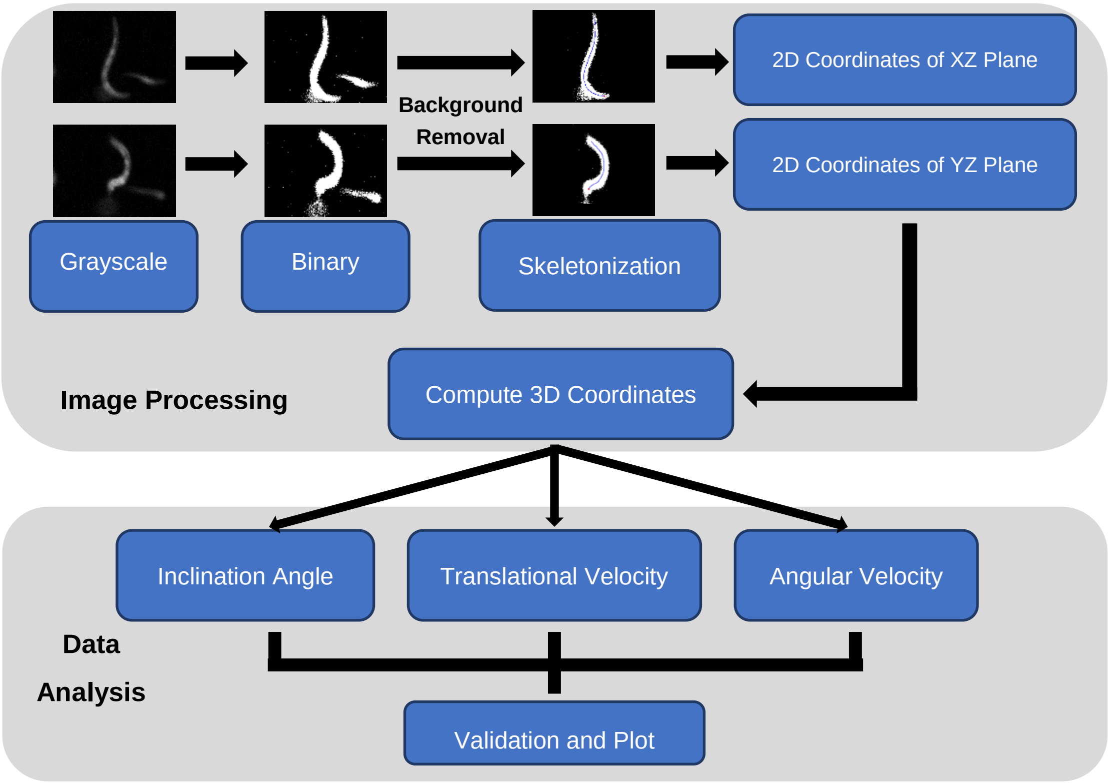


**Fig. S1: Schematic depiction of image processing and data analysis**. The X-Z and Y-Z projections of a worm were captured with cameras 1 and 2, respectively. The red dot and blue lines in the skeletonization step identify, respectively, the worm's head and skeleton.

6. The worm's skeleton is defined as the center line of the worm's body and found by thinning the worm's body from both sides simultaneously. The end points of this skeleton were defined as the worm's head and tail (The head was distinguished from the tail manually in the first image.), and the angle formed by the line connecting the head and the tail relative to the direction opposite to the gravity vector was defined as the angle of inclination θ (**Fig. 1**).

7. Once the positions of the end points were determined in the two focal planes, 3D

coordinates were computed for the head and tail.

While we minimized the possibility of two or more worms crossing each other's paths using a dilute worm suspension (1.5 worms/μL), images in which worm paths overlapped, as assessed by visual inspection, were censored. We estimate that less than 15 frames of all frames analyzed were censored.

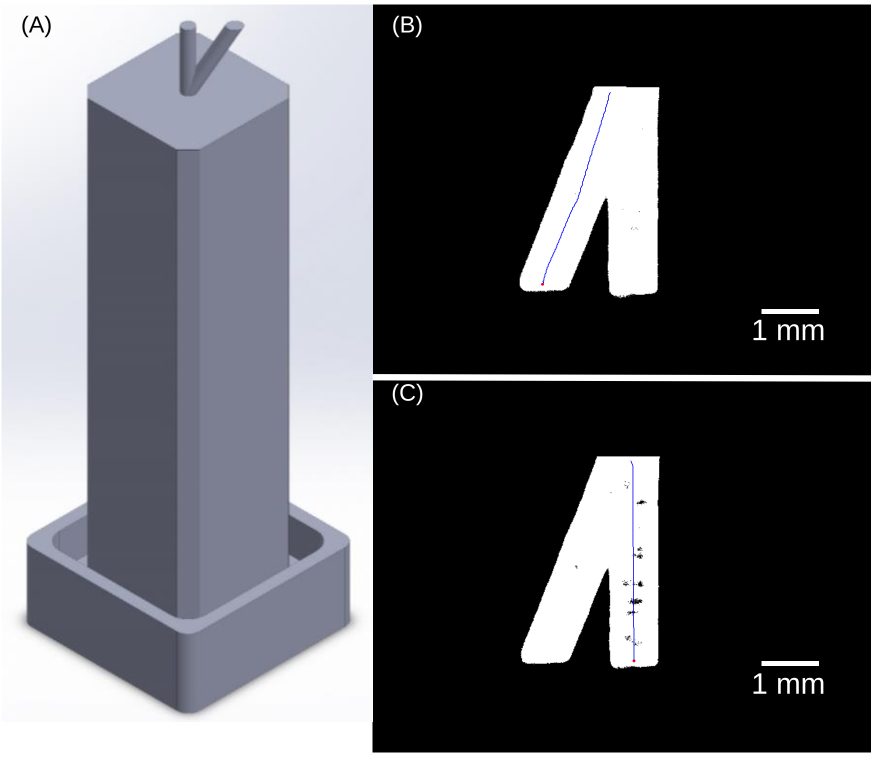


**Fig. S2: Calibration jig**. (A) The isometric view of the 3D printed calibration jig. Our calculations were verified by reproducing the inclination angle of the jig's legs: $\theta = 150°$ and $\varphi = 45°$ (A) and $\theta = 180°$ (B).

### Image Processing Verification

To verify our analysis, we 3D-printed (Formlabs, Form 2) a calibration jig (**Fig. S2**). The jig includes a square base sized to fit tightly into the cuvette, a vertical leg ($\theta = 180°$) located at the jig base's center, and an inclined leg ($\theta = 150°$, $\varphi = 45°$).

We calculated the angle of the inclined leg as $\theta=149.6° \pm 2.1°$ and the vertical leg $\theta=179.81° \pm 1.07°$. We estimate that our calculation of the inclination angle (equation S2) is accurate within 3.5°. The cameras are aligned within 7.39 $\mu m$ in the vertical direction.

### S2. Reproducibility and Potential Artifacts

We repeated the experiments on different days with similar results. **Fig. S3** depicts the polar angle θ kernel (probability density) function estimate at two different days showing similar results.

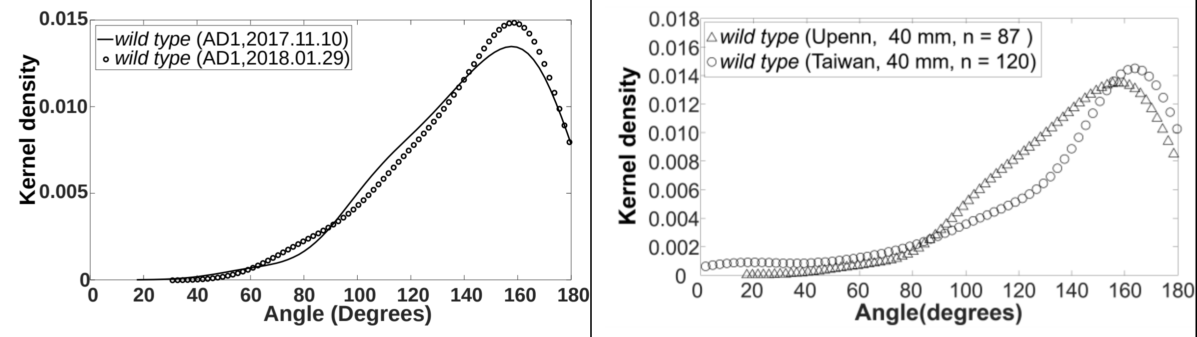


**Fig. S3**: **Polar angle kernel (probability density) function estimate obtained in two experiments carried out 10 weeks apart (left) and in two different continents (USA and Taiwan).**

To test whether our data may have been biased by convective currents in the experimental apparatus, we examined the azimuthal angle φ kernel (probability) density function (**Fig. S4**). These data show that worms descended with a nearly uniformly distributed azimuthal angle.

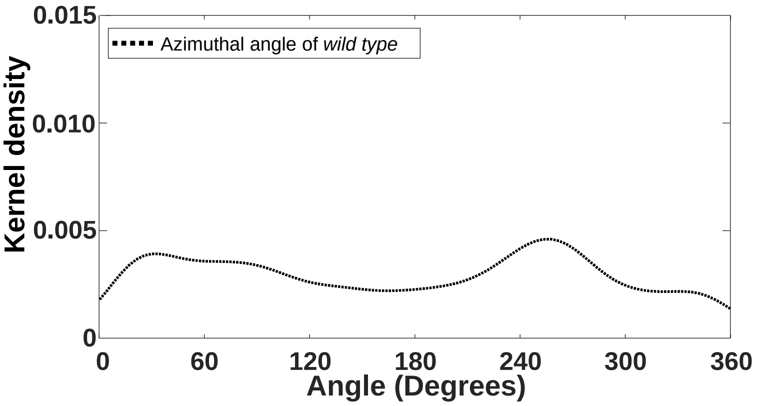


**Fig. S4**: **Azimuthal angle $\varphi$ kernel (probability) density estimate.**

### S3. Kernel Distribution Estimate (KDE)

To construct the KDE and the corresponding Cumulative Distribution Function (CDF) from experimental data, we used standard Matlab commands with default values, including the default, optimized bandwidth $h$ and gaussian distribution for the kernel function. To examine the effect of the bandwidth on the KDE, we repeated the calculations with various bandwidths for the case of day-one adult wild type animals (**Fig. S5**). **Fig. S5** demonstrates that KDE produces similar results for a broad selection of bend widths. Our results therefore are unlikely to be biased by our choice of the bandwidth h.

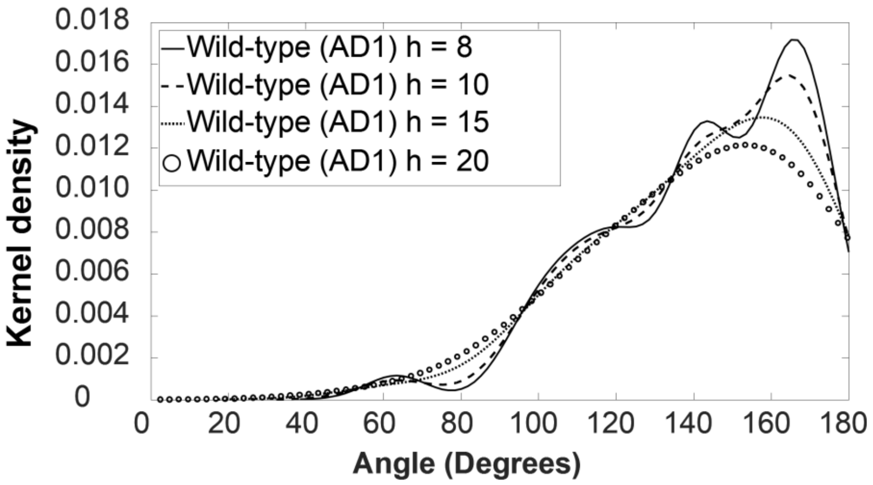


**Fig. S5**: **KDE with various selections of bandwidth (h).** Wild type animals in water.

### S4. Directional Statistics

Often directional distributions on a unit sphere is modeled with von Mises-Fisher (vMF) probability distribution function

$$f(\theta,\varphi)=\frac{\lambda}{4\pi Sinh\lambda}e^{\lambda\cos(\pi-\theta)}. \qquad \text{(S1)}$$

In the above, λ is the *concentration parameter* (a reciprocal measure of dispersion). Since our data suggests that the distribution is independent of the azimuthal angle φ, we integrated (S1) accounting for the spherical geometry to obtain

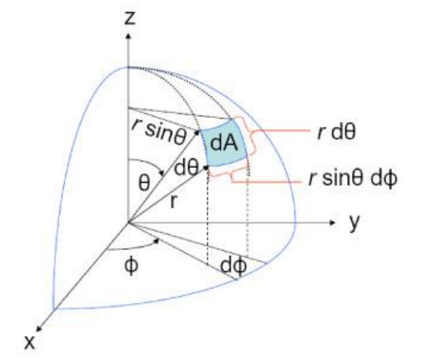


**Fig. S6**: **An area element on a sphere.** ([https://math.stackexchange.com/questions/131735/surface-element-in-spherical-coordinates](https://math.stackexchange.com/questions/131735/surface-element-in-spherical-coordinates) )

$$f(\theta)=\int_0^{2\pi} f(\theta,\varphi)\sin\theta d\varphi=\frac{\lambda}{2Sinh\lambda}e^{\lambda\cos(\pi-\theta)}\sin(\pi-\theta). \qquad \text{(S2)}$$

$\int_0^{\pi} f(\theta)d\theta=1.$ λ →0 corresponds to a uniform distribution:

$$\lim_{\lambda->0} f(\theta)=\lim_{\lambda->0}\frac{\lambda}{2Sinh\lambda}e^{\lambda\cos(\pi-\theta)}\sin(\pi-\theta)=\frac{1}{2}\sin(\pi-\theta). \qquad \text{(S3)}$$

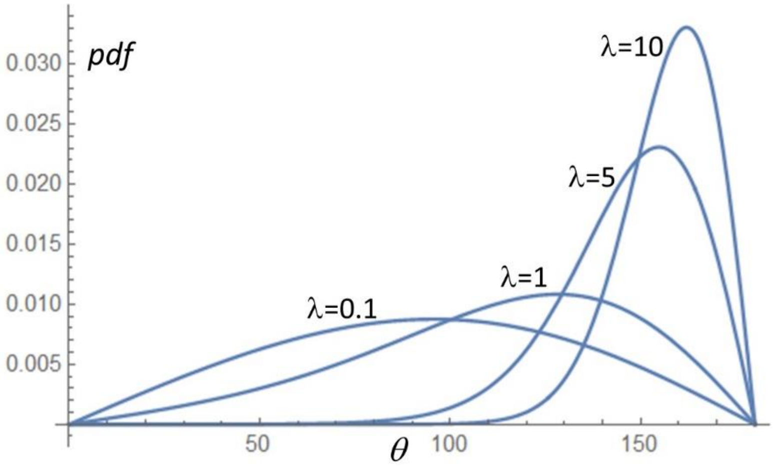


**Fig. S7:** Von Mises - Fisher probability distribution function for various values of the concentration factor λ = 0.1, 1, 5, and 10.

Fig. S7 Depicts the probability distribution function (pdf) *f(θ)* as a function of *θ* when λ=0.1, 1, 5, and 10. The pdf for λ=0.1 closely resembles a uniform distribution (sin function).

The Cumulative distribution function is (CDF)

$$CDF\left(\theta\right)=\frac{1}{2\,\mathrm{Sinh}\,\lambda}\left(e^{-\lambda\cos(\theta)}-e^{-\lambda}\right) \quad \text{(S4)}$$

**Fig. S8** depicts the cumulative distribution function (cdf) as a function of $\theta$ when λ=0.1, 1, 5, and 10.

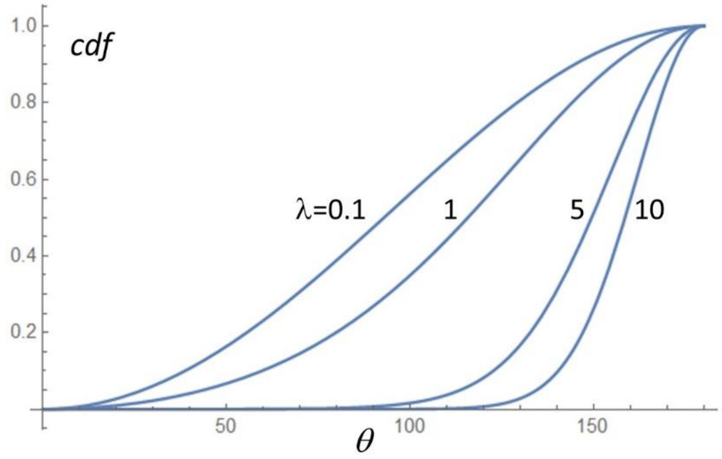


**Fig. S8:** Von Mises - Fisher cumulative distribution function for various values of the concentration factor λ = 0.1, 1, 5, and 10.

We obtained the experimental estimate of λ by fitting equation (S4) to our experimental CDF. When λ = 1.73 of the animals exhibited a polar angle θ > 90°. This fraction increased as λ increased. We consider cases with $\lambda > 1$ as exhibiting gravitaxis.

**S5. Kernel distribution estimate of well-fed wild type animals at various depths under the liquid surface**

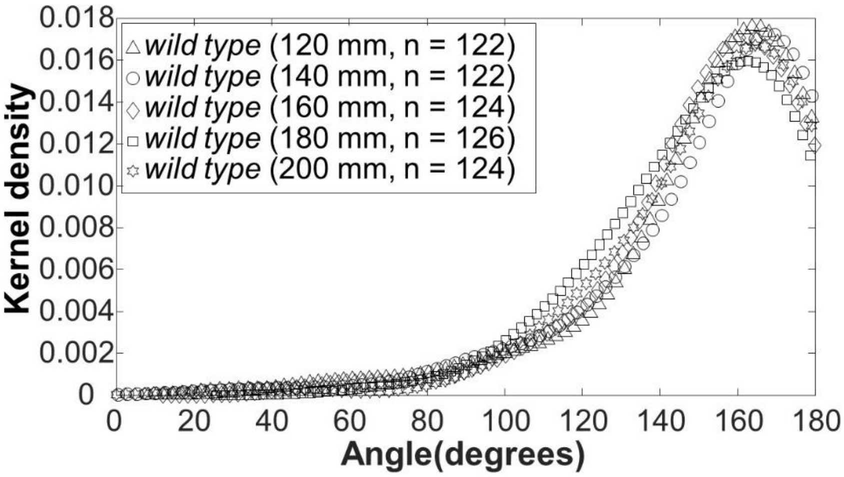


**Fig. S9**: **Wild-type worms retain their orientation after a certain residence time**. Kernel-density of wild-type swimmers' orientation angle (θ) at positions 120, 140, 160, 180, and 200 mm beneath the liquid surface.

**S6. Paralyzed animals do not align with the direction of gravity**

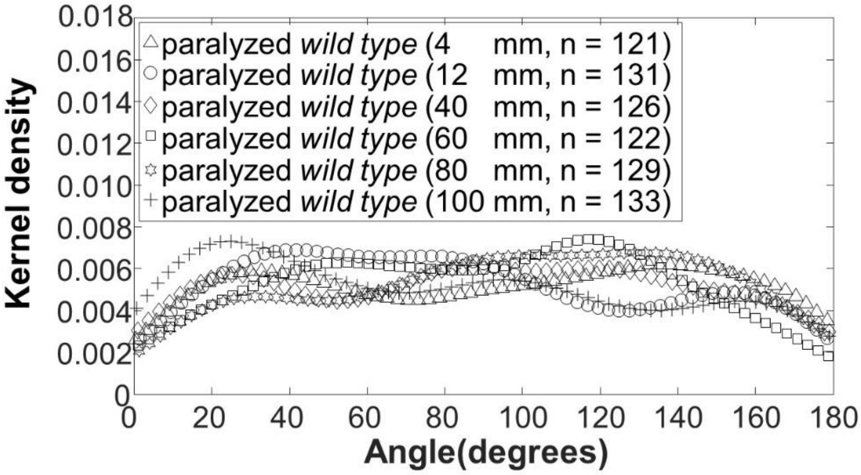


**Fig. S10**: **Paralyzed WT worms retain random distribution of their orientation as they settle in solution.** Kernel (probability) density estimate (KDE) of heat-shocked paralyzed WT animals at depths ranging from 4 mm to 100 mm beneath the liquid surface. The bandwidth of the KDE smoothing window is 15. This figure augments the data presented in the main text (**Fig. 5**) for longer residence times. **Fig. 5** and this figure demonstrates that orientational KDE of paralyzed WT animals does not vary with depth and residence time.

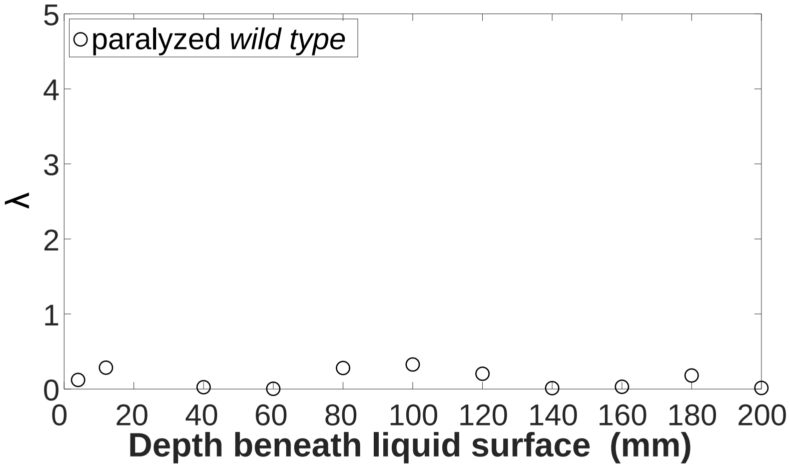


**Fig. S11**: **Paralyzed WT animals: concentration parameter λ as a function of depth (residence time).** λ remains close to zero, independent of depth (and hence independent of residence time in solution), indicating that the animal's orientation is randomly distributed.

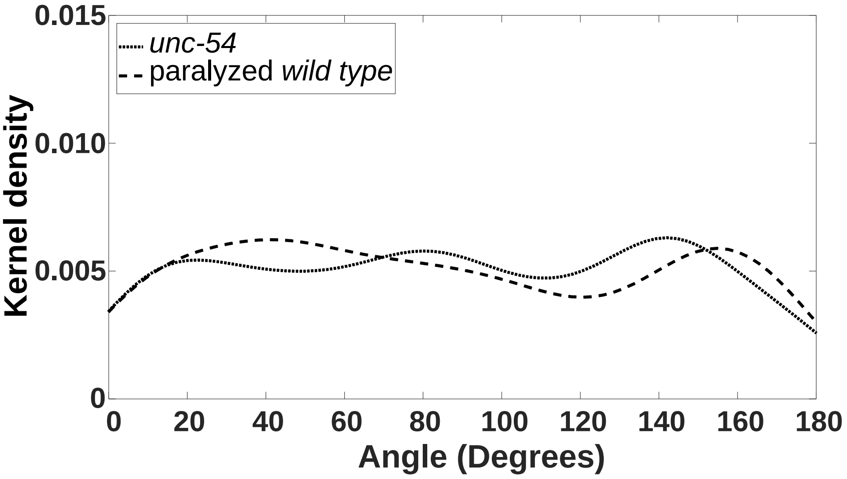


**Fig. S12**: **KDE of unc-54 is compared with that of paralyzed WT at d = 40 mm.** The two KDEs are similar and consistent with a random distribution in orientation angle.

### S7: Starved WT and motion-impaired animals (*unc-29*) align with the gravitational field at a lower rate than well-nourished WT animals

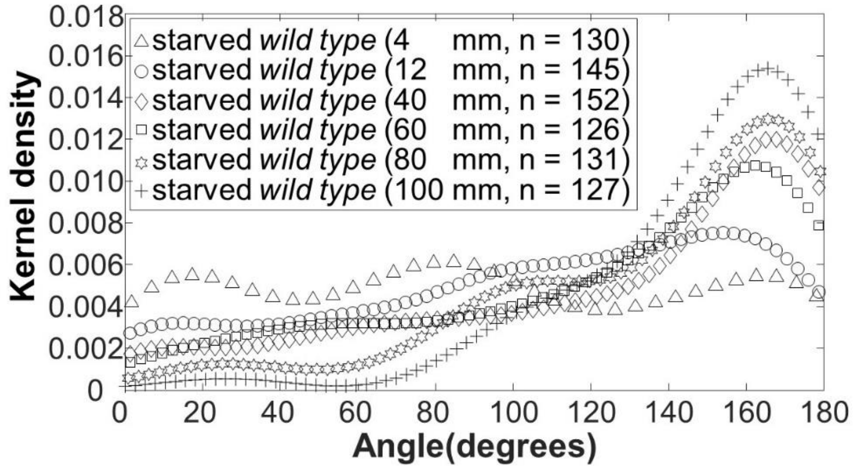


**Fig. S13**: **Starved (>1 h after last feeding) WT animals align with the direction of the gravity vector.** Kernel (probability) density estimate (KDE) of starved WT animals at depths ranging from 4 to 100 mm beneath the liquid surface. The bandwidth of the KDE smoothing window is 15.

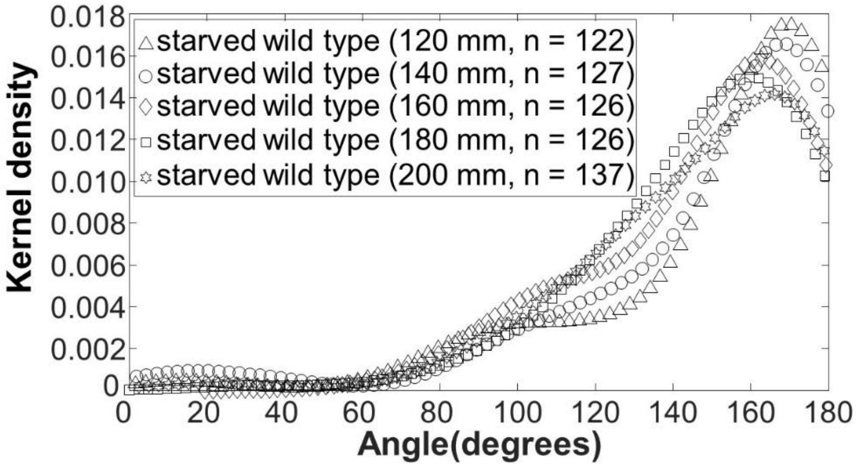


**Fig. S14**: **Starved (>1 h after last feeding) WT animals align with the direction of the gravity vector.** Kernel (probability) density estimate (KDE) of starved WT animals at depths ranging from 120 to 200 mm beneath the liquid surface. The bandwidth of the KDE smoothing window is 15.

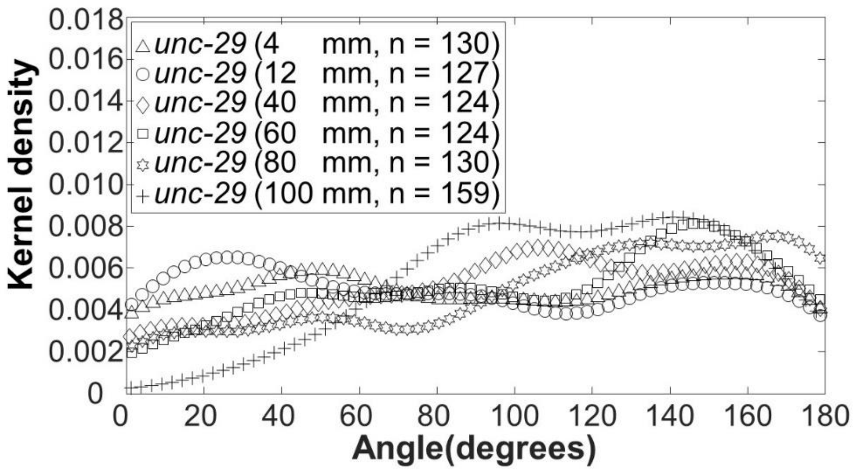


**Fig. S15**: **Animals defective in muscle function** (*unc-29*) **align slowly with the direction of the gravity vector.** Kernel (probability) density estimate (KDE) *unc-29* mutants at depths ranging from 4 to 100 mm beneath the liquid surface. The bandwidth of the KDE smoothing window is 15.

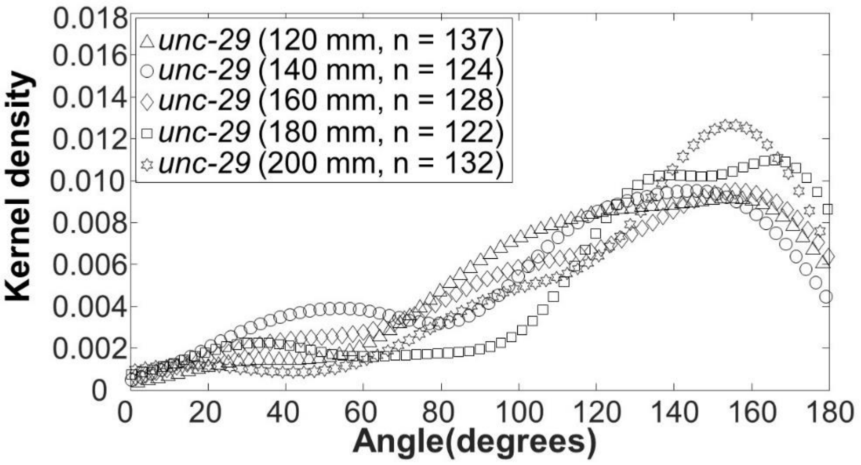


**Fig. S16**: **Animals defective in muscle function** (*unc-29*) **align with the direction of the gravity vector.** Kernel (probability) density estimate (KDE) *unc-29* mutants at depths ranging from 120 to 200 mm beneath the liquid surface. The bandwidth of the KDE smoothing window is 15.

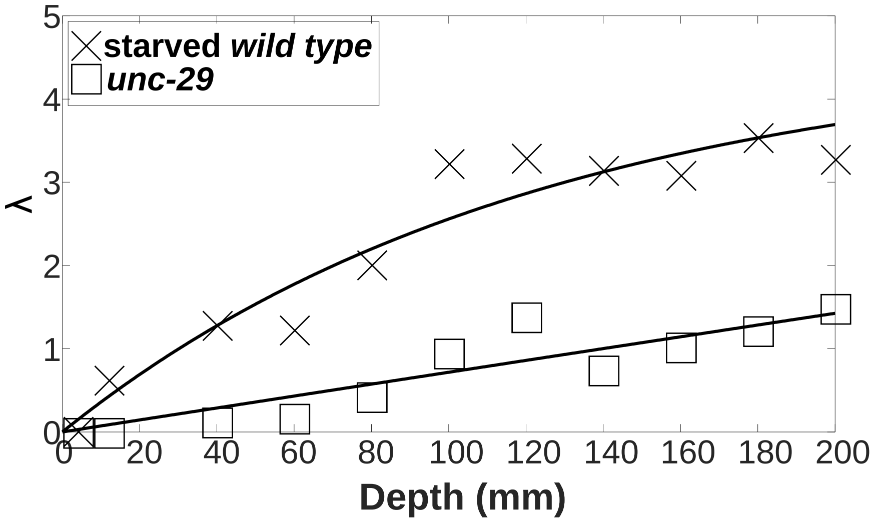


**Fig. S17**: **The concentration factor of starved WT animals and *unc-29* mutants increases as their submersion depth (residence time) increases.** The increase in the concentration factor indicates alignment with the direction of the gravity vector. The solid lines are best fit curves (equation 3) with $\lambda_\infty \sim 4.60$ and $\beta \sim 0.008\ mm^{-1}$ (starved WT) and $\lambda_\infty \sim 5.9$ and $\beta \sim 0.001\ mm^{-1}$ (*unc-29*).

**S8. Well-fed adult worms of the AB1 Adelaide strain align their direction of swimming with the direction of the gravity vector**

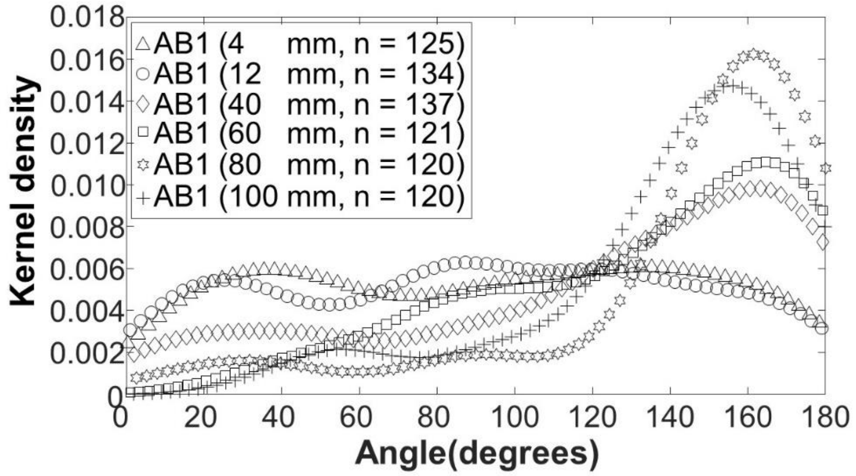


**Fig. S18**: **Well-fed adult worms of the AB1 Adelaide strain align their direction of swimming with the direction of the gravity vector.** Kernel (probability) density estimate AB1 mutants at depths ranging from 4 to 100 mm beneath the liquid surface. The bandwidth of the KDE smoothing window is 15.

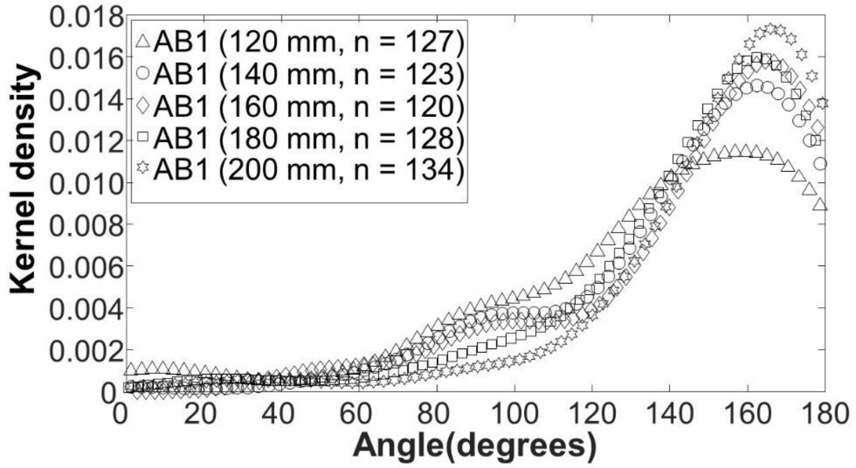


**Fig. S19**: **Well-fed adult worms of the AB1 Adelaide strain align their direction of swimming with the direction of the gravity vector.** Kernel (probability) density estimate AB1 mutants at depths ranging from 120 to 200 mm beneath the liquid surface. The bandwidth of the KDE smoothing window is 15.

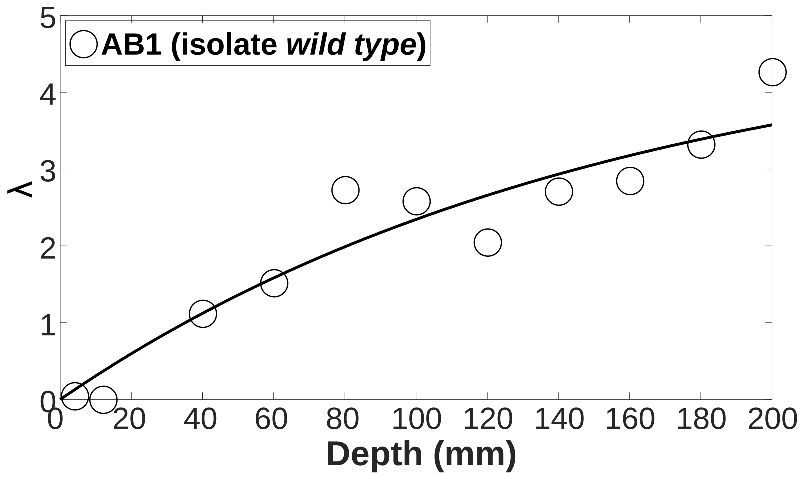


**Fig. S20: The concentration factor of well-fed AB1 animals increases as their submersion depth (residence time) increases.** The increase in the concentration factor indicates alignment with the direction of the gravity vector. The solid line is a best fit curve (equation 3) with $\lambda_\infty \sim 4.93$ and $\beta \sim 0.009\ mm^{-1}$.
